# Effect of differences in mechanical stress in vivo on the onset and progression of knee osteoarthritis

**DOI:** 10.1101/2021.06.04.447099

**Authors:** Kohei Arakawa, Kei Takahata, Yuichiro Oka, Kaichi Ozone, Sumika Nakagaki, Saaya Enomoto, Kenji Murata, Naohiko Kanemura, Takanori Kokubun

**Author notes:** **Corresponding Author:** Takanori Kokubun, PhD, 820 Sannomiya, Koshigaya-Shi, Saitama, 343-8540. Japan. Authors’ email addresses: KA,; KT,; YO,; KO,; SN,; SE,; KM,; NK,; TK.

## Abstract

**Objective:** The effect of the type of mechanical stress on OA onset has not been clarified. The aim of this study was to establish a new model that reproduces the type and increase and decrease of mechanical stress in vivo and to clarify the differences in the mechanism of knee OA onset and progression among the models.

**Design:** To reproduce the difference in mechanical stress, we used the anterior cruciate ligament transection (ACL-T) model and the destabilization of the medial meniscus (DMM) model. In addition, we created a controlled abnormal tibial translation (CATT) model and a controlled abnormal tibial rotation (CATR) model that suppressed the joint instability of the ACL-T and DMM model, respectively. These four models reproduced the increase and decrease in shear force due to joint instability and compressive stress due to meniscal dysfunction. We performed joint instability analysis with soft X-ray, micro computed tomography analysis, histological analysis, and immunohistological analysis in 4 and 6 weeks.

**Results:** Joint instability decreased in the CATT and CATR groups. The meniscus deviated in the DMM and CATR groups. Chondrocyte hypertrophy increased in the ACL-T and DMM groups with joint instability. In the subchondral bone, bone resorption was promoted in the ACL-T and CATT groups, and bone formation was promoted in the DMM and CATR groups.

**Conclusions:** Increased shear force causes articular cartilage degeneration and osteoclast activation in the subchondral bone. In contrast, increased compressive stress promotes bone formation in the subchondral bone earlier than articular cartilage degeneration occurs.

## INTRODUCTION

Osteoarthritis (OA) is a degenerative joint disease characterized by articular cartilage degeneration, osteophyte formation, and subchondral bone sclerosis^1,2^. Knee OA has the highest morbidity among OA, and interferes with daily life due to pain and dysfunction of the knee joint. The mechanism of knee OA onset remains unclear. Radin et al.^3^ had reported that the imbalance of subchondral bone strength causes articular cartilage degeneration; since then, it has been debated whether the onset of knee OA is caused by articular cartilage degeneration or subchondral bone changes. Among the studies that had used an OA animal model, some reports have shown that articular cartilage degeneration precedes changes in subchondral bone^4,5^. However, other reports suggest that changes in subchondral bone precede articular cartilage degeneration^6,7^. Since it is difficult to reproduce the complicated pathology of knee OA with a single model, it is necessary to evaluate the characteristics of each model^8^.

The mechanism of knee OA onset has not been elucidated, but mechanical stress is considered to be the main factor^9^. Although excessive mechanical stress on joints causes articular cartilage degeneration, moderate mechanical stress contributes to articular cartilage protection^10^. In addition, a study that had used an animal model of knee OA revealed that moderate exercise with a treadmill suppressed the progression of knee OA.

In contrast, high-intensity exercise promoted the progression of knee OA^11,12^. This suggests that mechanical stress acts on cartilage degeneration in knee OA in an intensity-dependent manner. Mechanical stress such as shearing force, compressive stress, and stretching force, are typically applied to the knee joint^13^. The impact of the intensity of these mechanical stresses are clearly defined, but their individual effects have not yet been clarified.

The rodent knee OA model mainly includes the anterior cruciate ligament transection (ACL-T) model and the destabilization of the medial meniscus (DMM) model. Both models were defined to induce knee OA by causing joint instability. Joint instability causes an increase in shear force on the articular cartilage of the knee. However, the direction and magnitude of joint instability differ between the two models^14^. Degeneration of the medial meniscus occurs rapidly in the DMM model^15^. The meniscus provides joint stability and reduces the compressive stress of the tibial plateau. Therefore, in the DMM model, it is inferred that the shear force increases due to rotational joint instability, and that the compressive stress increases due to meniscal dysfunction. The DMM model also causes milder joint instability and slower articular degeneration than those caused by the ACL-T model^14,16^. However, no reports have focused on the difference in mechanical stress between these two models.

We hypothesized that the type of mechanical stress determines the mechanism of knee OA onset. It is difficult to reproduce the types of mechanical stress applied to in vivo experiments because multiple types of mechanical stress are constantly applied to the joint. Therefore, in vitro verification is required to reproduce the types of mechanical stress. However, it has been reported that articular cartilage and subchondral bone interact in the knee joint in vivo^17^. Therefore, the effect of the type of mechanical stress on the onset and progression of cartilage degeneration in knee OA can be determined by reproducing the type of mechanical stress and its increase and decrease in vivo.

We used the ACL-T and DMM models to reproduce the types of mechanical stresses in vivo. However, as mentioned above, shear force and compressive stress are added to the DMM model. Therefore, we applied the recently reported controlled abnormal tibial translation (CATT) model, which suppresses joint instability in the anterior-posterior direction caused by the ACL-T model^18^, to the DMM model. The CATT model reduces the increase in shear force caused by ACL rupture by suppressing joint instability. Therefore, we established a new controlled abnormal tibial rotation (CATR) model to suppress joint instability in the DMM model. By comparing these four models, we conceived that the increase and decrease in shear force and compressive stress in the living body could be reproduced.

Therefore, the purpose of this study was to establish a new model that reproduces the type and increase and decrease of mechanical stress in vivo and to clarify the differences in the mechanism of knee OA onset and progression among the models.

## MATERIALS AND METHODS

### Animals and experimental design

This study was approved by the Animal Research Committee of Saitama Prefectural University (approval number: 2020-1), and the animals were handled in accordance with the relevant legislation and institutional guidelines for humane animal treatment. In this study, 59 adult (12-week-old) ICR (Institute for Cancer Research) male mice were randomized into one of five groups: ACL-T group (ACL-T, n = 12), CATT group (CATT, n = 12), DMM group (DMM, n = 12), CATR group (CATR, n = 12), and sham group (Sham, n=11). All mice were housed in plastic cages, and the room had a 12-hour light/dark cycle. Mice were permitted unrestricted movement within the cage and had free access to food and water.

### Surgical procedures

All surgical procedures were performed on the left knee joint with the mice under a combination anesthetic (medetomidine, 0.375 mg/kg; midazolam, 2.0 mg/kg; and butorphanol, 2.5 mg/kg). The medial capsule was exposed in the ACL-T group, and scissors were used to cut the ACL to generate knee OA. CATT was performed following the same procedure as the ACL-T surgery (Fig. 1A). Mice assigned to the CATT group had bone tunnels created in the distal femur and proximal tibia using a 25-gauge needle and 4-0 nylon threads threaded through them to suppress the anterior-posterior joint instability that occurs in the ACL-T model (Fig 1Ba,b). Subsequently, 4-0 nylon threads were tightly tied (Fig. 1Bc) to compensate for ACL function and suppress anterior-posterior joint instability.

**Figure 1.**
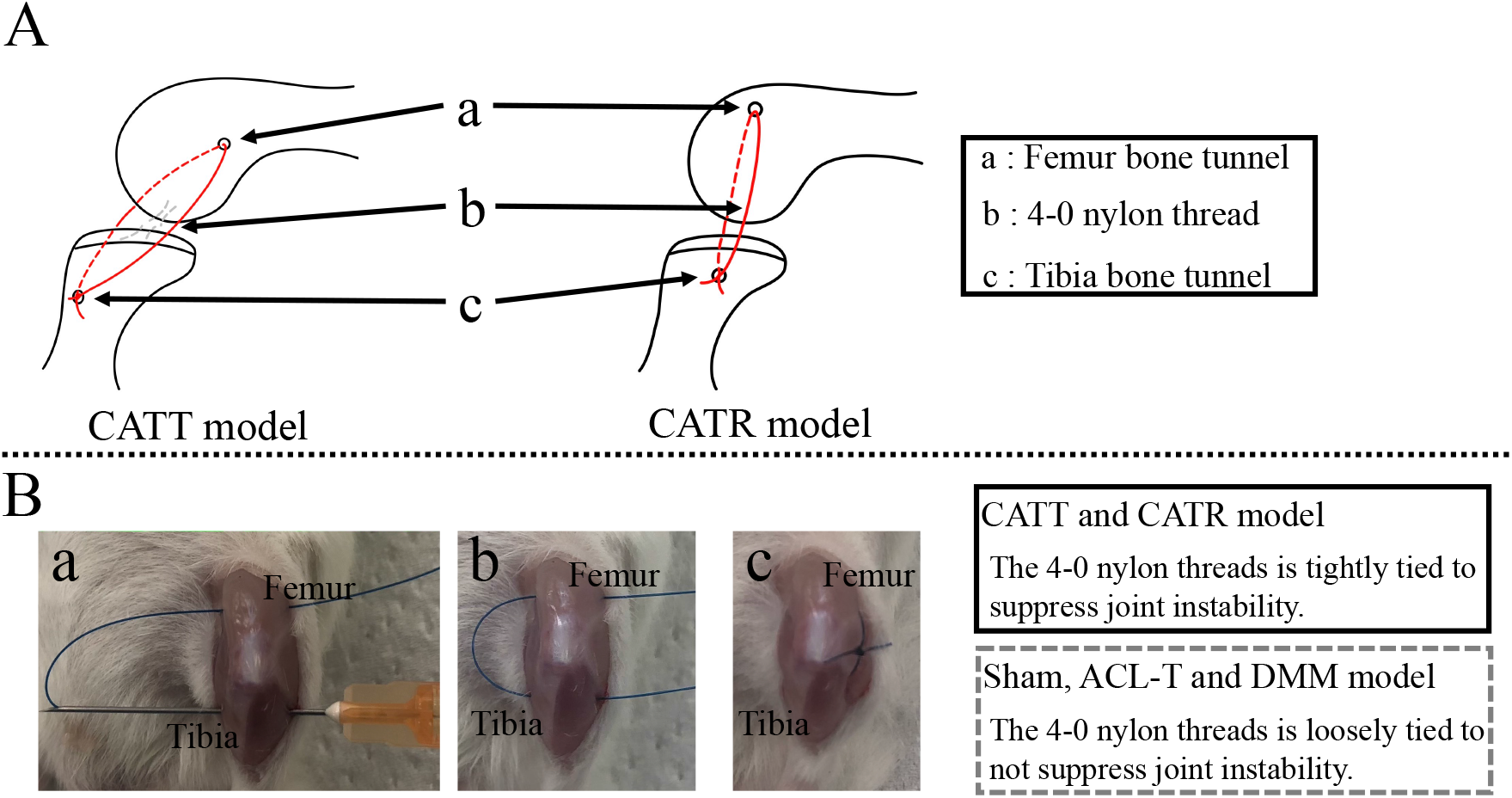
Surgery for the CATT and CATR model. (A) Schematic diagram of the CATT and CATR models. Nylon suture suppresses anterior-posterior joint instability caused by the ACL-T model and rotational joint instability caused by the DMM model. (B) Details of surgery. Create a bone tunnel distal to the femur and proximal to the tibia (a). Use 4-0 nylon suture to penetrate the bone tunnel (b). Tie nylon thread in a loop (d).

DMM surgery was performed on the left knee joint of the mice as previously described^16^, in which the medial capsule was exposed and scissors were used to cut the medial meniscotibial ligament (MMTL). CATR surgery was performed following the same procedure as DMM surgery (Fig. 1A). Mice assigned to the CATR group had bone tunnels created in the distal femur and proximal tibia using a 25-gauge needle and 4-0 nylon threads threaded through them to suppress the joint rotational instability that occurs in the DMM model (Fig. 1Ba,b). Subsequently, 4-0 nylon threads were tightly tied (Fig. 1Bc) to physically suppress joint rotational instability. To eliminate differences between the groups, bone tunnels were created in the Sham, ACL-T and DMM groups as well as in the CATT and CATR groups, and a nylon thread was tied loosely so as not to suppress joint instability.

### Anterior drawer test

To assess anterior-posterior joint instability, we performed the anterior drawer test using a constant force spring (0.05 kgf; Sanko Spring Co., Ltd., Fukuoka, Japan) and a soft X-ray device (M-60; Softex Co., Ltd., Kanagawa, Japan)^19^. At 4 and 6 weeks after surgery, we collected the knee joints of mice with an intact femur, tibia, and foot. Digital images were acquired using an x-ray sensor (Naomi; RF Co. Ltd., Nagano, Japan) with a voltage of 28 kV, a current of 1.5 mA, and an exposure time of 1 s. Soft x-ray images were used to quantify the anterior displacement of the tibia using a dedicated image analysis software (ImageJ; National Institutes of Health, Bethesda, MD, USA).

### Tibial rotational test

To assess rotational joint instability, we performed a tibial rotational test using a constant force spring (0.05 kgf; Sanko Spring Co., Ltd., Fukuoka, Japan) and a soft x-ray device (M-60; Softex Co., Ltd., Kanagawa, Japan). At 4 and 6 weeks after surgery, we collected the knee joints of mice with an intact femur, tibia, and foot. For this experiment, the femur and tibia were fixed with the knee joint at 90° of flexion. In addition, a 25-gauge needle was passed through the tibia as a measure of tibial rotation change. In the test, the proximal tibia was pulled laterally and medially with a 4-0 nylon thread. We photographed x-rays during medial and lateral traction and defined the angular change between the needle and the vertical line of the device as rotational instability. Digital images were acquired using an x-ray sensor (Naomi; RF Co. Ltd., Nagano, Japan) with the settings described above. Based on the soft x-ray image, changes in the tibial rotation angle were quantified using a dedicated image analysis software (ImageJ; National Institutes of Health, Bethesda, MD, USA).

### Analysis of the subchondral bone using micro-computed tomography (μCT)

For analysis of the subchondral bone, each knee joint was fixed with 4% paraformaldehyde. The knee joints were scanned using a *μ*CT system (Skyscan 1272, BRUKER, MA, USA) with the following parameters: pixel size, 6μm; voltage, 60 kV; current, 165 μA. Subsequently, the reconstructed image was acquired using the NRecon software (BRUKER, MA, USA). We designated the region of interest on 40 slides of the medial tibial subchondral bone. We then calculated the bone volume/tissue volume fraction (BV/TV, %), trabecular thickness (Tb.Th, mm), trabecular number (Tb.N, 1/mm), and trabecular separation (Tb.Sp, mm) using CTAn software (BRUKER, MA, USA). In addition, the position of the medial meniscus was macroscopically observed from the reconstructed image.

### Histological analysis

Mice were sacrificed at 4 and 6 weeks after surgery, and the knee joint was collected and fixed in 4% paraformaldehyde for 1 day, followed by decalcification in 10% ethylenediaminetetraacetic acid for 3 weeks, dehydrated in 70% and 100% ethanol and xylene, and embedded in paraffin blocks. The samples were cut in the sagittal plane (6 μm thickness) using a microtome (ROM-360; Yamato Kohki Industrial Co., Ltd., Saitama, Japan). Safranin-O/fast green staining was performed to evaluate the degree of articular cartilage degeneration. The Osteoarthritis Research Society International (OARSI) histopathological grading system was used to assess cartilage degeneration for structural changes and fibrillation lesions^20^ by two independent observers (K Takahata and SN) who were blinded to all other sample information. The average value of the scores of the two observers was used as the representative value.

### Immunohistochemical Analysis and TRAP Staining

To evaluate the expression of type X collagen and Osterix, we performed immunohistochemical staining using the avidin-biotinylated enzyme complex method and the VECTASTAIN Elite ABC Rabbit IgG Kit (Vector Laboratories, Burlingame, CA, USA). The tissue sections were deparaffinized with xylene and ethanol, and antigen activation was performed using proteinase K (Worthington Biochemical Co., Lakewood, NJ, USA) for 15 min. Endogenous peroxidase was inactivated with 0.3% H2O2/methanol for 30 min. Nonspecific binding of the primary antibody was blocked using normal goat serum for 30 min. The sections were then incubated with anti-collagen X antibody (1:300, ab58632, Abcam) and Osterix polyclonal antibody (1:250, bs-1110R, Bioss) overnight at 4°C. Afterward, the sections were incubated with biotinylated secondary anti-rabbit IgG antibody and stained using Dako Liquid DAB+ Substrate Chromogen System (Dako, Glostrup, Denmark).

Cell nuclei were stained with hematoxylin. For analysis, we calculated the ratio between the number of Type Ⅹ collagen positive cells and the number of chondrocytes in an articular cartilage area of 10000 μm^2^ (100 μm × 100 μm). In the analysis of Osterix, the positive cell rates in the medullary cavity of the subchondral bone at 2500 μm^2^ (50 μm × 50 μm) were compared. We calculated the Osterix positive cell rate at two locations, and the average value was used as the representative value. According to manufacturer instructions, osteoclast activity was detected by histochemical staining for tartrate-resistant acid phosphatase (TRAP) using the TRAP stain kit (Wako, Osaka, Japan). We then calculated the osteoclast surface (Oc.S/BS, %) in the bone marrow region as an analysis of osteoclast activity.

### Statistical analysis

All data were analyzed using R software, version 3.6.1. First, the normality of all data was verified using the Shapiro-Wilk test. Since the evaluation of joint instability and the OARSI score were nonparametric, the Kruskal-Wallis test was used. Subsequently, the Steel-Dwass test was used for post-hoc analysis. A one-way analysis of variance (ANOVA) was performed for the other data, followed by a post hoc Tukey-Kramer test. Parametric data were expressed as means with 95% confidence intervals, whereas non-parametric data were expressed as medians with interquartile ranges. The statistical significance was set at p<0.05.

## RESULTS

### Evaluation of joint instability

Anterior-posterior joint instability was quantified in terms of anterior displacement of the tibia using the anterior drawer test (Fig. 2A). At 4 weeks, the anterior displacement of the ACL-T group increased significantly compared to that of the other groups. The anterior displacement of the CATT group significantly reduced the anterior-posterior joint instability of the ACL-T group, but significantly increased it compared to the Sham, DMM, and CATR groups. ([ACL-T vs. Sham], p=0.031; [ACL-T vs. CATT], p=0.032; [ACL-T vs. DMM], p=0.031; [ACL-T vs. CATR], p=0.031; [CATT vs. Sham], p=0.032; [CATT vs. DMM], p=0.032; [CATT vs. CATR], p=0.032).

**Figure 2.**
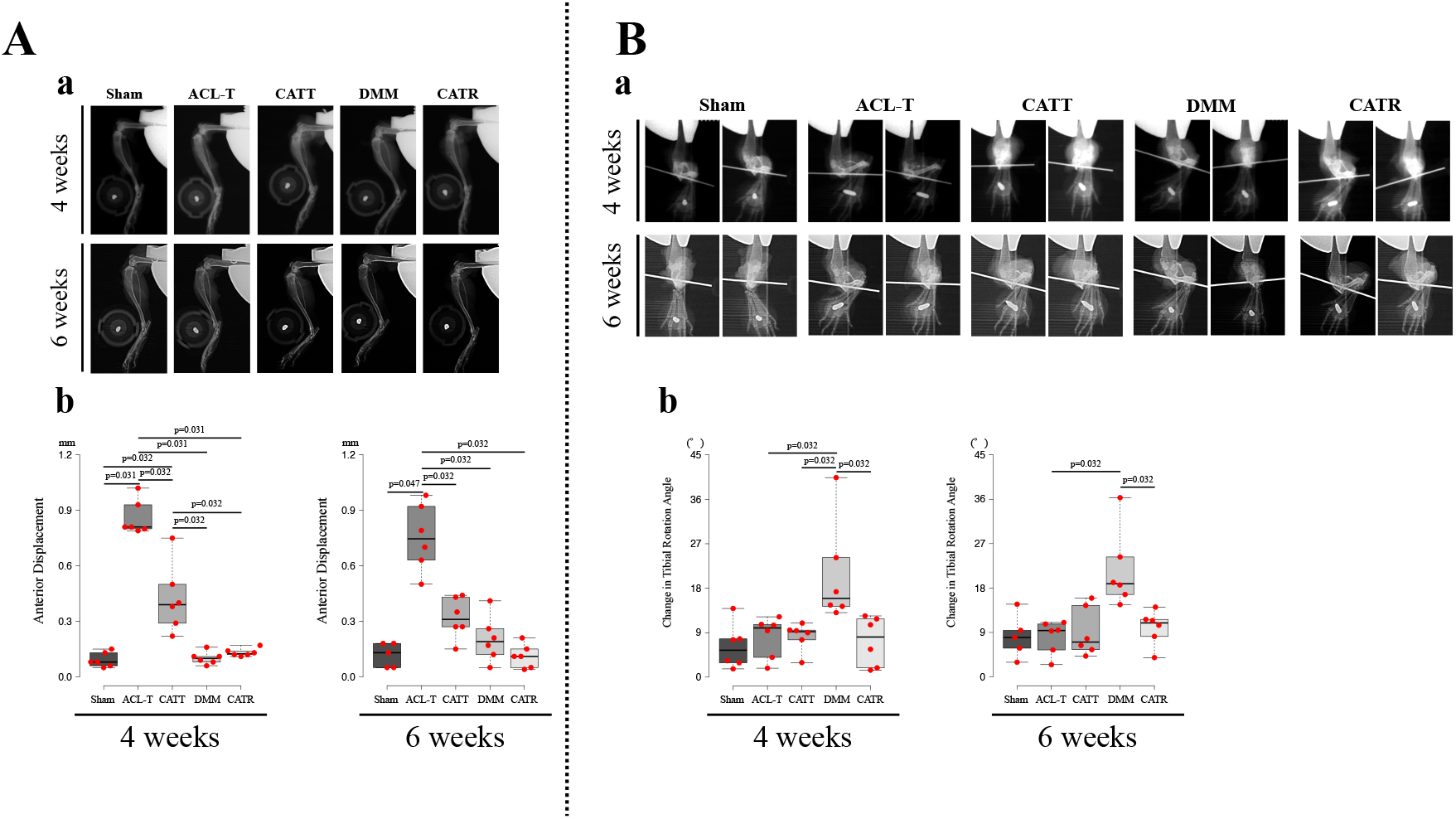
Evaluation of joint instability using soft x-ray analysis. (A) Results of the anterior drawer test. Anterior tibial displacement was observed in the ACL-T group (a). At both 4 and 6 weeks, the amount of anterior tibial displacement in the ACL-T group was increased compared to that in other groups. The CATT group suppressed the joint instability that occurred in the ACL-T group (b). The data are presented as the median with interquartile range. (B) Results of the tibial rotational test. An increase in tibial rotation angle was observed in the DMM group (a). At both 4 and 6 weeks, the amount of change in tibial rotation angle in the CATR group was smaller than that in the DMM group. The CATR group suppressed the joint instability that occurred in the DMM group (b). The data are presented as the median with interquartile range.

Even at 6 weeks, the anterior displacement of the ACL-T group increased significantly compared to that of the other groups. On the other hand, the CATT group significantly reduced the anterior displacement of the ACL-T group, and no significant difference was observed compared to the other groups ([ACL-T vs. Sham], p =0.047; [ACL-T vs. CATT], p =0.032; [ACL-T vs. DMM], p =0.032; [ACL-T vs. CATR], p =0.032).

Rotational joint instability was quantified in terms of anterior displacement of the tibia using a tibial rotational test (Fig. 2B). At 4 weeks, the change in tibial rotation angle in the DMM group were not significantly different from those in the Sham group but were significantly increased compared to that of the ACL-T, CATT, and CATR groups. In contrast, the change in the tibial rotation angle in the CATR group was not significantly different from that in the Sham, ACL-T, or CATT groups ([DMM vs. ACL-T], p =0.032; [DMM vs. CATT], p=0.032; [DMM vs. CATR], p=0.032).

Similarly, at 6 weeks, the change in tibial rotation angle in the DMM group were not significantly different from that in the Sham group but were significantly increased compared to that of the ACL-T and CATR groups. In contrast, the change in the tibial rotation angle in the CATR group was not significantly different from that in the Sham, ACL-T, or CATT groups ([DMM vs. ACL-T], p =0.032; [DMM vs. CATR], p =0.032).

### Subchondral bone changes between models

The 3D reconstruction images of the knee joint by μCT analysis are shown in Fig. 3A. At 4 and 6 weeks, lateral deviations of the medial meniscus were confirmed in the DMM and CATR groups, but no deviation of the medial meniscus was observed in the other groups.

**Figure 3.**
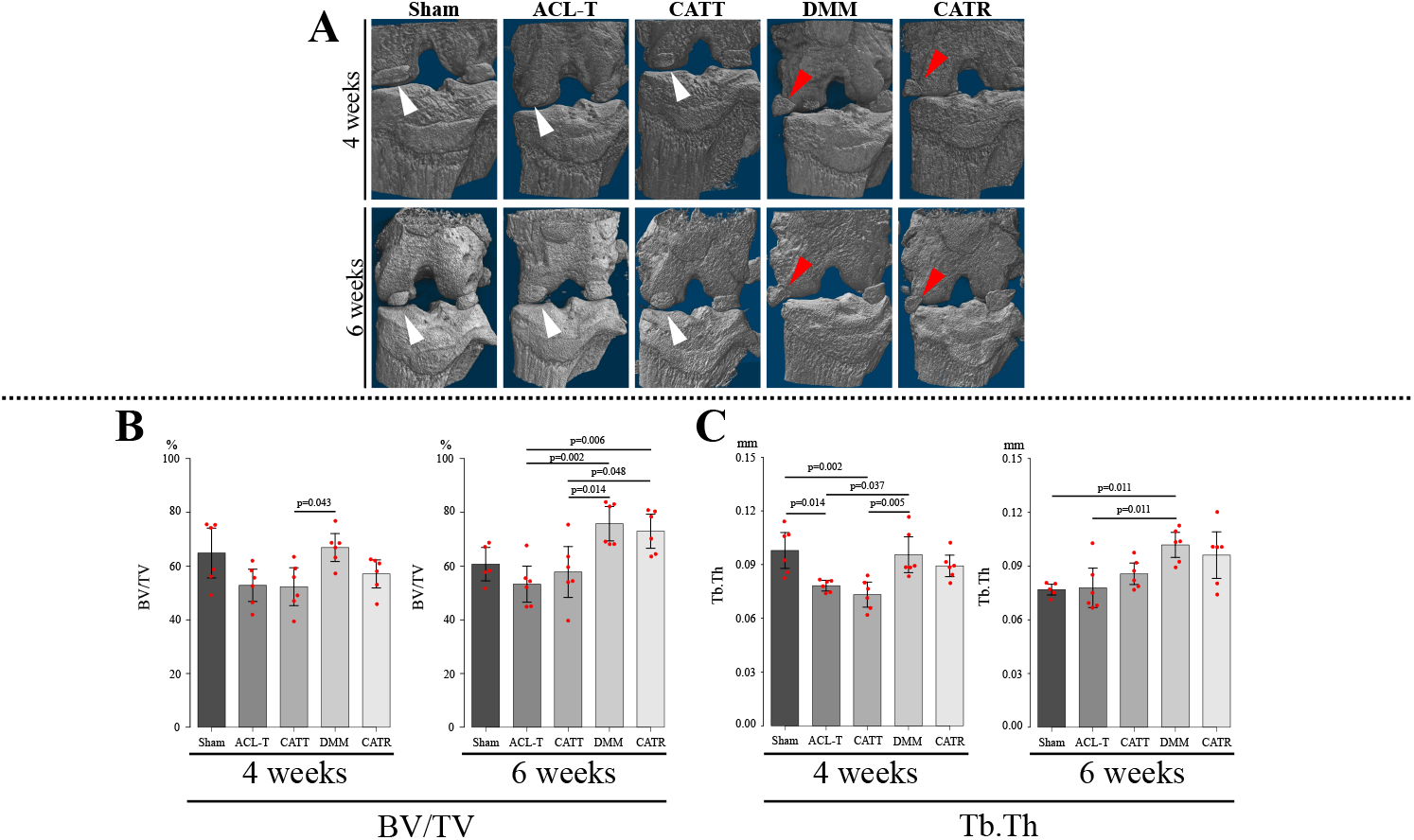
Analysis result by μCT. (A) Comparison of lateral displacement of the medial meniscus. Lateral displacement of the medial meniscus was observed in the DMM and CATR groups (red triangle). No displacement of the medial meniscus was observed in the Sham, ACL-T, and CATT groups (white triangle). (B) Results of BV/TV. At 4 weeks, the DMM group showed higher values than the CATT group, and at 6 weeks, the DMM and CATR groups showed higher values than the ACL-T and CATT groups. Data are presented as the mean with 95% CI. (C) Results of Tb.Th. At 4 weeks, the ACL-T and CATT groups showed lower values than the Sham and DMM groups. At 6 weeks, the DMM group showed higher values than the Sham and ACL-T groups. Data are presented as the mean with 95% CI.

The results of the BV/TV in each group are shown in Fig. 3B. At 4 weeks, the BV/TV in the DMM group was significantly higher than that in the CATT group. (p=0.043). Although there was no significant difference, the BV/TV in the ACL-T and CATT groups was lower than that in the Sham group, and the BV/TV in the DMM group was higher than that in the Sham group (Fig. 3B). At 6 weeks, the BV/TV in the DMM and CATR groups was significantly higher than that in the ACL-T and CATT groups ([DMM vs. ACL-T], p=0.002; [DMM vs. CATT], p=0.014; [CATR vs. ACL-T], p=0.006; [CATR vs. CATT], p=0.048). Although there was no significant difference, the BV/TV in the ACL-T and CATT groups was lower than that in the Sham group, and the BV/TV in the DMM and CATR groups were higher than those in the Sham group.

The Tb.Th results are shown in Fig. 3C. At 4 weeks, Tb.Th in the ACL-T and CATT groups were significantly lower than that in the Sham and DMM groups ([ACL-T vs. sham], p=0.014; [ACL-T vs. DMM], p=0.037; [CATT vs. Sham], p=0.002; [CATT vs. DMM], p=0.005). At 6 weeks, Tb.Th in the DMM group was significantly higher than that in the sham and ACL-T groups ([DMM vs. Sham], p=0.011; [DMM vs. ACL-T], p =0.011). Although there was no significant difference, Tb.Th in the CATR group was higher than that in the Sham group. There was no significant difference in Tb.N and Tb.Sp between the groups at 4 and 6 weeks (data not shown).

### Increased joint instability promotes superficial articular cartilage degeneration

The results of safranin-O/fast green staining are shown in Fig. 4A. Surface fibrillation of the articular cartilage was observed in the ACL-T group at 4 and 6 weeks. Although there was no significant difference, the OARSI score of the ACL-T group was higher than that of the other groups at 4 weeks (Fig. 4B). Similarly, there was no significant difference in OARSI score at 6 weeks (Fig. 4B).

**Figure 4.**
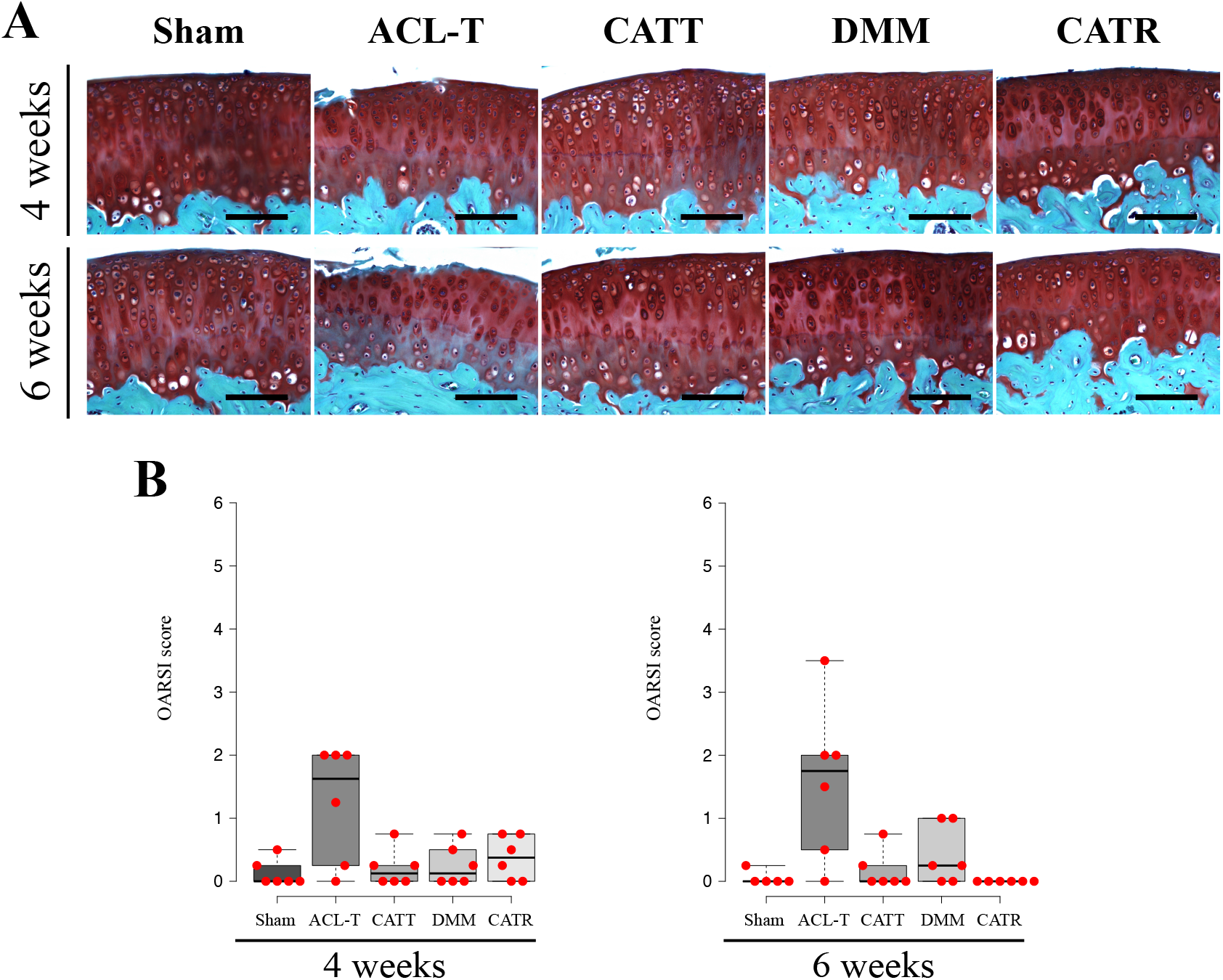
Comparison of articular cartilage degeneration by OARSI score. (A) Representative safranin-O/fast green-stained image of each group. In the ACL-T group, irregularity and fibrillation of articular cartilage were observed. (B) The results of OARSI score. There was no significant difference in OARSI scores. The data are presented as the median with interquartile range. Scale bar: 100 μm.

The results of immunohistochemical staining of Col X are shown in Fig. 5A. At 4 weeks, the positive cell rate of Col X in the ACL-T and DMM groups was higher than that in the Sham group (Fig. 5B) ([ACL-T vs. Sham], p <0.001; [DMM vs. Sham], p =0.002). At 6 weeks, the Col X-positive cell rate in the ACL-T and DMM groups was higher than that in the Sham group, and that in the CATR group was significantly lower than that in the ACL-T group (Fig. 5B) ([ACL-T vs. Sham], p=0.002; [DMM vs. Sham], p=0.022, [CATR vs. ACL-T], p=0.008).

**Figure 5.**
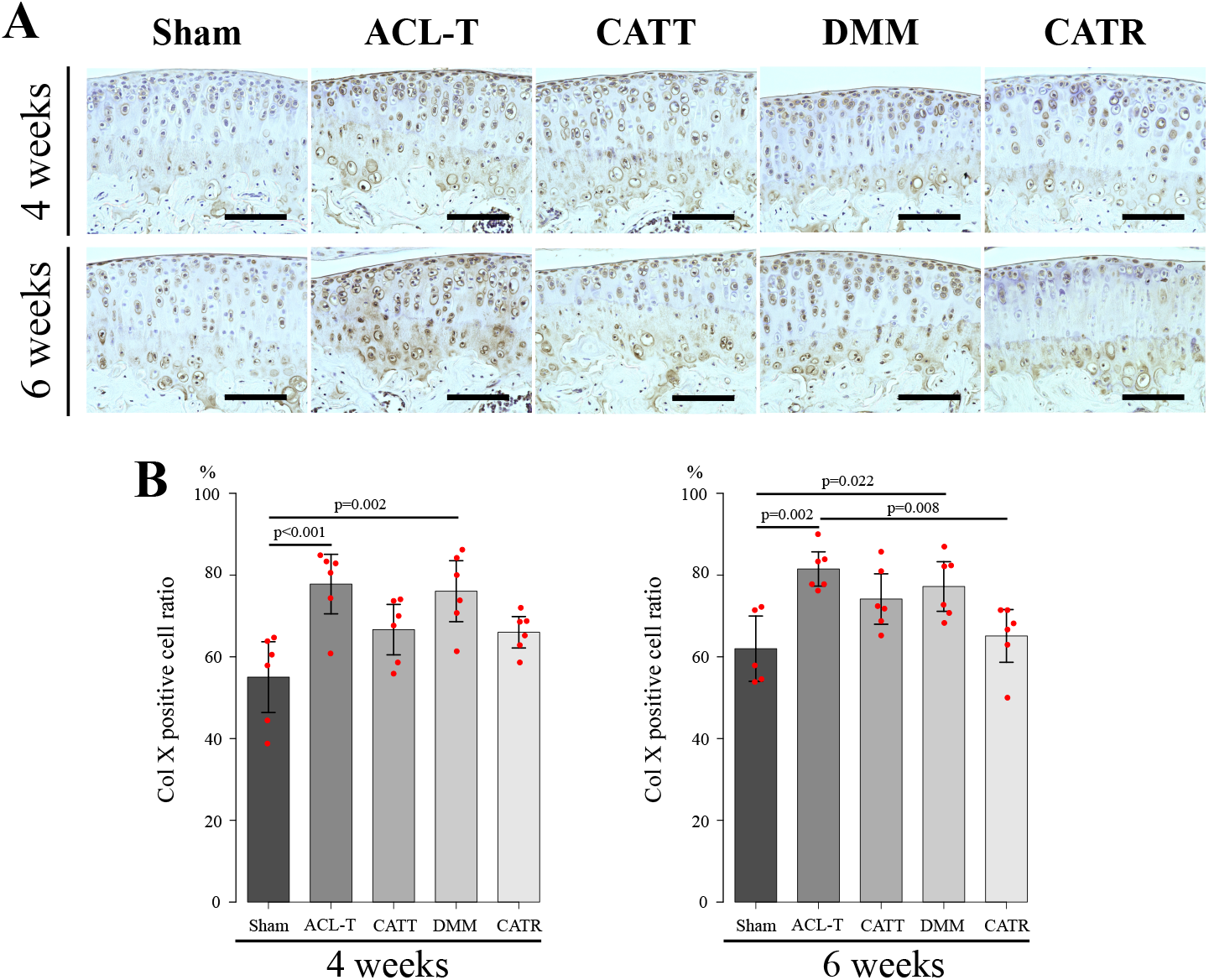
Comparison of Col X positive cell rates. (A) Representative Col X-stained image of each group. (B) At 4 weeks, the positive cell rate of the ACL-T and DMM groups was higher than that of the Sham group. Similarly, at 6 weeks, the positive cell rate of the ACL-T and DMM groups was higher than that of the Sham group. The positive cell rate in the CATR group was lower than that in the ACL-T group. Data are presented as the mean with 95% CI. Scale bar: 100 μm.

### Subchondral bone remodeling differences between models

The results of TRAP staining are shown in Fig. 6A. Regarding the activity of osteoclasts in subchondral bone, the Oc.S/BS in the ACL-T group was higher than that in the other groups at 4 weeks (Fig. 6B) ([ACL-T vs. Sham], p <0.001; [ACL-T vs. CATT], p =0.015, [ACL-T vs. DMM], p=0.001; [ACL-T vs. CATR], p=0.001). However, there was no significant difference in Oc.S/BS between the groups at 6 weeks(Fig. 6B).

**Figure 6.**
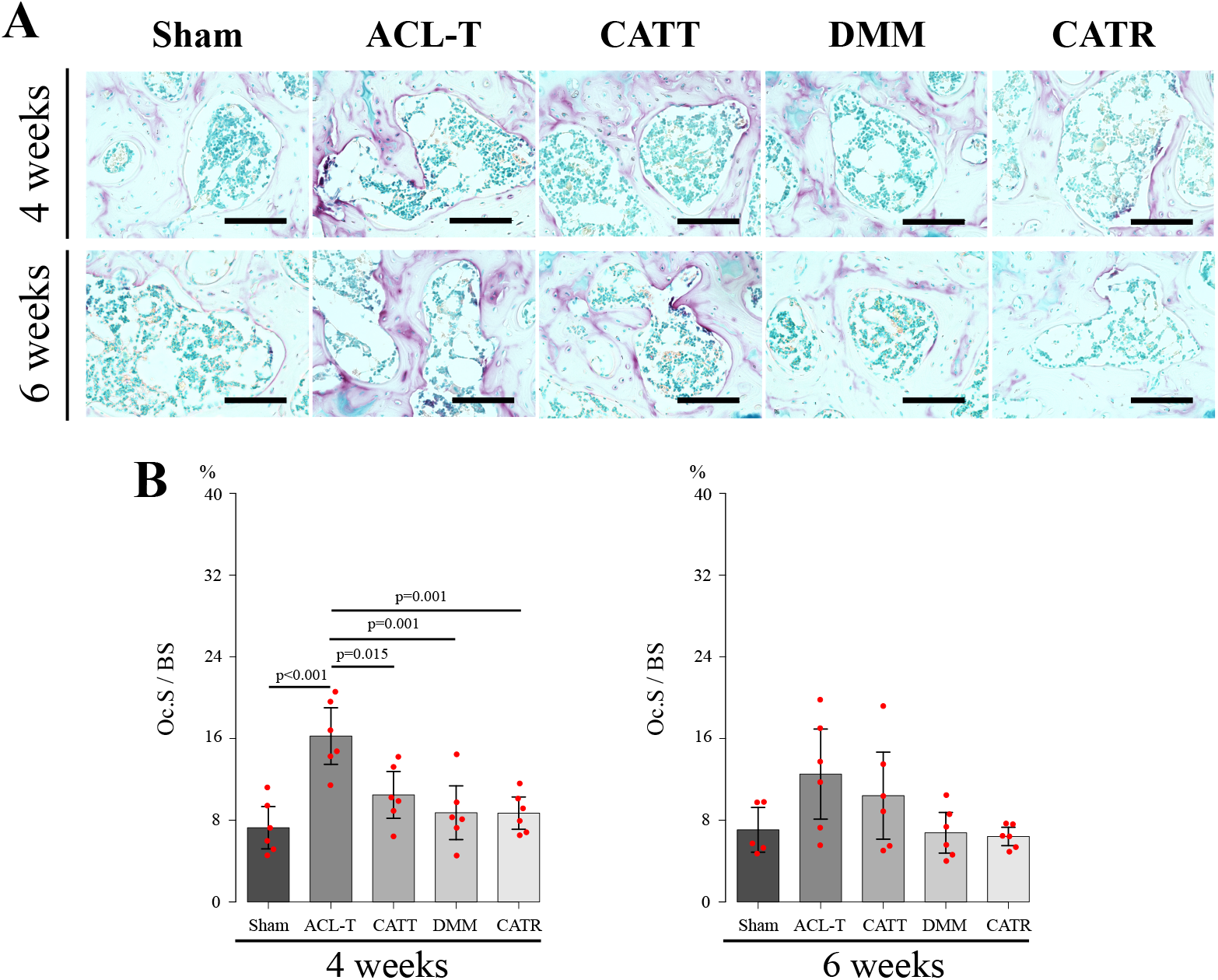
The results of TRAP staining. (A) Representative TRAP-stained image of each group. (B) The results of Oc.S/BS. At 4 weeks, Oc.S/BS in the ACL-T group was higher than that in the other groups. No significant difference was observed at 6 weeks. Data are presented as the mean with 95% CI. Scale bar: 100 μm.

The results of immunohistochemical staining of Osterix are shown in Fig. 7A. At 4 weeks, the positive cell rate of Osterix in the DMM and CATR groups was higher than that in the Sham group, and that in the DMM group was significantly lower than that in the ACL-T group (Fig. 7B) ([DMM vs. Sham], p =0.011; [CATR vs. Sham], p =0.035; [DMM vs. ACL-T], p =0.031). However, at 6 weeks, there was no significant difference in the positive cell rate of Osterix between the groups (Fig. 7B).

**Figure 7.**
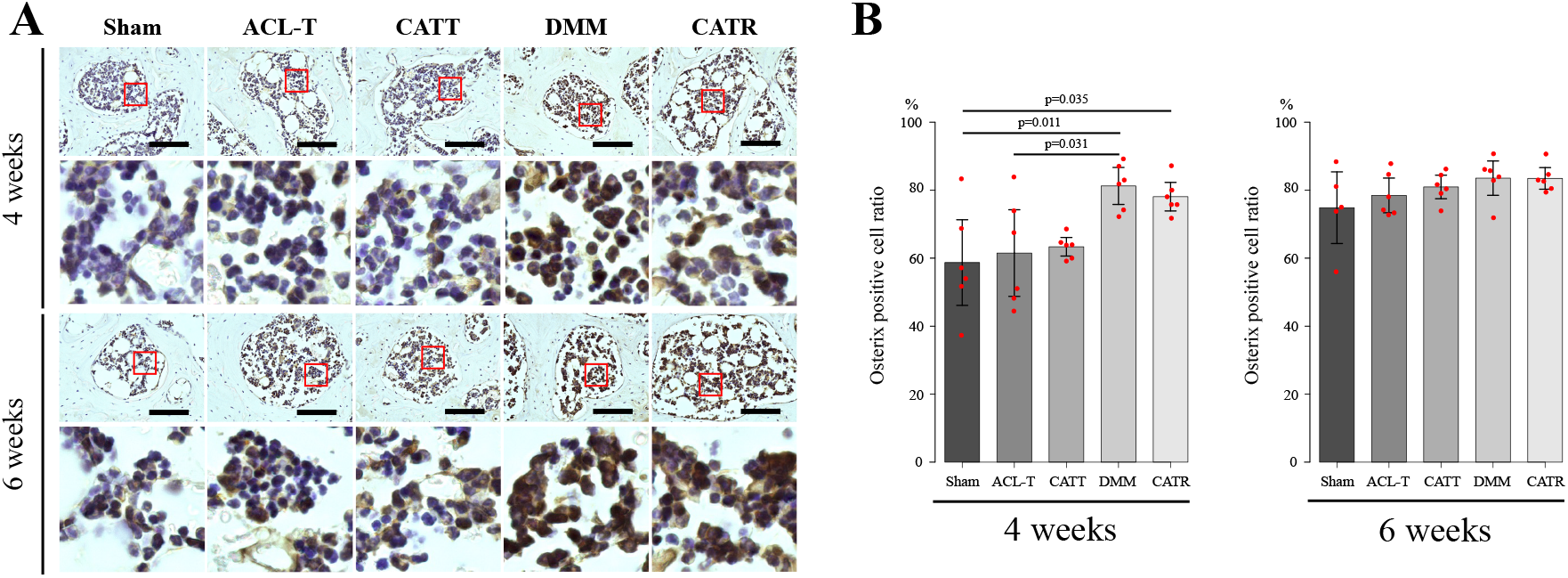
The results of immunohistochemical staining of Osterix. (A) Representative Osterix-stained image of each group. (B) Analysis results of Osterix positive cell rate. Analysis results of positive cell rate. At 4 weeks, the Osterix positive cell rate in the DMM group was higher than that in the Sham and ACL-T groups, and the positive cell rate in the CATR group was higher than that in the Sham group. There was no significant difference between the groups at 6 weeks. Data are presented as the mean with 95% CI. Scale bar: 100 μm.

## DISCUSSION

This study examined the differences in articular cartilage degeneration and subchondral bone changes in four different models of joint instability and meniscal dysfunction. CATT and CATR models have been established to control joint instability. The increase and decrease in shear force due to joint instability and compressive stress due to meniscal dysfunction were reproduced. Increased shear force causes articular cartilage degeneration and bone resorption of the subchondral bone, whereas increased compressive stress causes bone formation in the subchondral bone.

In this study, ACL-T and DMM models with joint instability and CATT and CATR models with suppressed joint instability were used. Soft x-ray analysis showed that the CATT group suppressed the joint instability produced by the ACL-T group (Fig. 2), and that the CATR group suppressed joint instability produced by the DMM group (Fig. 2). In our study, joint instability was defined as the shear force. Therefore, our results suggest that the shear force increased in the ACL-T and DMM groups, which increased joint instability, whereas the CATT and CATR groups, which suppressed joint instability, reduced the increase in shear force. In addition, lateral deviation of the medial meniscus was observed in the DMM and CATR groups. These results suggest that the CATR group suppresses joint instability but increases lateral deviation of the medial meniscus and meniscal dysfunction remains. In addition to contributing to joint stability, the meniscus disperses the compressive stress applied to the knee joint. Consequently, it can be inferred that compressive stress due to meniscal dysfunction increased in both the DMM and CATR groups.

It has been reported that articular cartilage degeneration depends on the magnitude of joint instability^21^. Adebayo et al.^14^ and Glasson et al.^16^ had reported that the DMM model has milder joint instability and slower articular cartilage degeneration than the ACL-T model. In addition, our colleague reported that articular cartilage degeneration is delayed by suppressing joint instability that occurs in the ACL-T model.^22–24^ Out histological results confirmed loss of the articular cartilage surface layer in the ACL-T group at 4 and 6 weeks. There was no significant difference in the OARSI score, but the CATT, DMM, and CATR groups had mild articular cartilage degeneration. The chondrocyte hypertrophy characteristic of early articular cartilage degeneration was significantly increased in the ACL-T and DMM groups, compared to the Sham group at 4 and 6 weeks. However, there was no significant difference between the CATT and CATR groups compared to the Sham group. These results support those of previous studies^21,22^ which suggest that increased joint instability causes articular cartilage degeneration. Our comparison of different models suggested that increased shear force due to the magnitude of joint instability was involved in superficial articular cartilage degeneration.

Subchondral bone changes are important for the onset and progression of OA. Abnormal subchondral bone remodeling is essential for OA pathology^14,25,26^. Previous reports that had induced subchondral bone remodeling in a single model reported that the ACL-T model causes bone loss^25^ and that the DMM model increases BV/TV early after the intervention^26^. We investigated the characteristics of subchondral bone changes by comparing four models with different mechanical stresses in vivo. Interestingly, μCT analysis showed two different characteristics of subchondral bone changes. The BV/TV decreased in the ACL-T and CATT groups with ACL disconnection. Nomura et al.^27^ had reported that reducing the mechanical stress on joints reduced the BV/TV. Therefore, we speculated that in the ACL-T and CATT groups, joint instability in the anterior-posterior direction caused the dispersion of compressive force and decreased BV/TV. To support these changes, osteoclast activity by TRAP staining was significantly increased in the Oc.S/BS in the ACL-T group. Therefore, we inferred that the dispersion of compressive stress contributes to the activation of osteoclasts and promotes bone resorption. In contrast, BV/TV increased in the DMM and CATR groups with dysfunction of the medial meniscus. BV/TV and Tb.Th increase when an axial compression load is applied to the tibia^28–30^. Thus, we speculated that in the DMM and CATR groups, the increased compressive force due to meniscal dysfunction promoted bone formation in the subchondral bone. To support these changes, the Osterix-positive cell rate was significantly higher in the DMM and CATR groups. Therefore, we inferred that the increase in compressive stress promotes the differentiation of osteoblasts and contributes to bone formation.

Summarize, the increase in shear force caused by joint instability resulted in articular cartilage degeneration and chondrocyte hypertrophy in articular cartilage and accelerated bone resorption in the subchondral bone. The increase in compressive stress due to meniscal dysfunction promoted bone formation in the subchondral bone earlier than the onset of articular cartilage degeneration. These results suggest that different types of mechanical stress in vivo may affect articular cartilage and subchondral bone differently in the development and progression of knee OA.

The main limitation of this study is that the types of mechanical stress could not be quantified. The type and increase or decrease in mechanical stress were reproduced from the difference in the functions of the ACL and meniscus. Therefore, it was necessary to verify the effect of different types of mechanical stress on the articular cartilage and subchondral bone using in vitro verification and a model that quantitatively applies a load to the knee joint. Therefore, the difference in subchondral bone changes observed in this study may be because the OA progression rate differs between models. Thus, future studies should investigate the difference in the OA progression mechanism of each model by performing longer-term verification and comparing the subchondral bone changes at the time of articular cartilage degeneration in each model.

In conclusion, our results suggest that increased shear force causes articular cartilage degeneration and osteoclast activation in the subchondral bone. In contrast, increased compressive stress promotes bone formation in the subchondral bone before articular cartilage degeneration occurs.

## Acknowledgements

The author(s) received no financial support for the research, authorship, and/or publication of this article. And we would like to thank Editage (http://www.editage.com) for English language editing.

## Author contributions

All authors approved the final submitted manuscript.

Study design: KA, KM, NK, and TK

Data collection, Histological analysis: KA, KT, YO, KO, SN and SE

Manuscript composition KA, TK

